# Characterizing Cone Spectral Classification by Optoretinography

**DOI:** 10.1101/2022.09.07.507027

**Authors:** Vimal Prabhu Pandiyan, Sierra Schleufer, Emily Slezak, James Fong, Rishi Upadhyay, Austin Roorda, Ren Ng, Ramkumar Sabesan

## Abstract

Light propagation in photoreceptor outer segments is affected by photopigment absorption and the phototransduction amplification cascade. Photopigment absorption has been studied using retinal densitometry, while recently, optoretinography (ORG) has provided an avenue to probe changes in outer segment optical path length due to phototransduction. With adaptive optics (AO), both densitometry and ORG have been used for cone spectral classification, based on the differential bleaching signatures of the three cone types. Here, we characterize cone classification by ORG, implemented in an AO line-scan OCT and compare it against densitometry. The cone mosaics of five color normal subjects were classified using ORG showing high probability (∼0.99), low error (<0.22%), high test-retest reliability (∼97%) and short imaging durations (< 1 hour). Of these, the cone spectral assignments in two subjects were compared against AOSLO densitometry. High agreement (mean: 91%) was observed between the two modalities in these 2 subjects, with measurements conducted 6-7 years apart. Overall, ORG benefits from higher sensitivity and dynamic range to probe cone photopigments compared to densitometry, and thus provides greater fidelity for cone spectral classification.

## 1. Introduction

Signals from the three cone spectral classes in the retina form the basis for color perception[1]. Their relative proportion and arrangement shapes and constrains how wavelength information from the external world is coded in the visual system to create the rich palette of hues that most humans perceive. Short-wavelength cones (S-cones) can be differentiated in histology [2], but the remarkably similar morphology and protein structure of long and middle-wavelength cones (LM-cones) has long precluded the creation of imaging techniques and antibodies to segregate them in a cellular-scale assay *ex vivo*. Until the advent of adaptive optics (AO) to overcome the optical aberrations of the eye [3] and resolve cone photoreceptors in living humans, there was virtually no information about the arrangement of individual L and M-cones in primate retina. In fact, AO retinal densitometry to delineate the spectral types of cones by Roorda & Williams [4] was one of the very first scientific applications of AO upon its introduction to the eye in 1997.

Densitometry has long been used to study various biophysical properties in cone photoreceptors, including waveguiding efficiency, the optical density and photosensitivity of chromophores housed in their outer segments[5-8]. Its principle rests on using the absorption of light by photopigments as a gauge to measure its properties. For cone spectral typing, differential absorption by the three photopigments in L, M and S-cones is used to separate them by their spectral type. An AO fundus camera was first used for this task, enabling the visualization of cone spectral types in humans and non-human primates [4, 9, 10]. In 2015, the feasibility of using an AO scanning laser ophthalmoscope (AOSLO) to classify cone spectral types was tested[11]. In addition to the advantages of increased axial and lateral resolution, contrast and signal-to-noise ratio, AOSLO densitometry allowed imaging the dynamic changes in photoreceptors upon light capture. Together, this enabled cone spectral classification with high efficiency with the time taken per subject ranging between 3 to 9 hours, and requiring an average of 8 – 15 repeat measurements per bleaching condition to boost the signal-to-noise ratio. In contrast, the AO fundus camera needed averages of up to 50 repeat measurements across 5 days in each bleach condition[4].

Recently, Zhang et al. 2019, 2021[12, 13] showed improved precision and efficiency for mapping cone spectral types using an AO optical coherence tomography (AO-OCT) instrument in phase-resolved configuration. The principle of the measurements rests on imaging light-evoked changes in the backscattered optical phase arising in the cone outer segment in response to a bleaching stimulus, also referred to as optoretinography or ORG [14, 15]. These changes in phase were converted to optical path length (ΔOPL) to serve as a quantifiable gauge for measuring light propagation after bleaching in the outer segment and the magnitude of ΔOPL scaled with increasing bleach strength. By adopting a bleaching light that differentially activated L, M and S-cones, it was demonstrated that the ΔOPL response arising from individual cones could be readily segregated into three cone sub-types. Following the general protocol established by Zhang et al. 2019, Pandiyan et al. 2020, 2021[16, 17] demonstrated the feasibility of cone spectral classification using a high-speed line-scan AO-OCT. Notably, cone spectral types were classified at an eccentricity of ∼0.3 deg. from the foveola enabling the first in vivo demonstration of decreasing S-cone proportion in the fovea[17].

It is instructive to compare the two paradigms in terms of their mechanism of action and dynamic range. As explained below, ORG has greater than 20x greater dynamic range for comparing photopigments across cone spectral types as compared to densitometry. This provides a fundamental benefit to the signal-to-noise ratio in ORG for classification; signal corresponds to a direct measure of photopigment concentration while noise is contributed by various common factors for both including low retinal reflectivity, the limit to light collection set by the numerical aperture of the eye, eye movements, reflections from retinal layers besides the cones, blood flow and more. The higher dynamic range is expected to reach similar or better accuracy in ORG with lower numbers of repeat measures, thus improving speed & efficiency.

Densitometry uses the same light source for imaging photoreceptors and bleaching photopigment, typically centered at 550nm where the spectral sensitivity of L and M-cones is high, *and* similar to each other. After dark-adaptation and regeneration of cone photopigment, the retina is exposed to the imaging & bleaching light. The backscattered light encodes the cone locations in the retinal image, and simultaneously carries information about absorption by the photopigment. For instance, the S-cones are relatively insensitive to a 550nm imaging light and therefore maintain their backscattered intensity after exposure to it. L and M-cone photopigments on the other hand absorb this light and appear darker than the S-cones immediately after light onset. With increasing bleach of L&M-cone photopigments, their concentration, and hence absorption decreases, consequently leading to an increase in the backscattered image intensity. This relationship between image intensity and photopigment concentration for L & M-cones is depicted in Figure 1a (brown & black dashed curves respectively). The change in image intensity immediately after stimulus onset and after a full bleach is referred to as the optical density and signifies the maximum photopigment-dependent signal available for probing its characteristics. Neitz et al. 2020[18] measured individual cone optical density using AOSLO densitometry for a range of visible wavelengths from 496nm to 598nm. A maximum optical density of 0.41 log10 units (2.6 times) was observed at 543 nm, i.e., a 2.6x change in image intensity is available as the dynamic range to probe LMS-cone photopigments.

**Figure 1:**
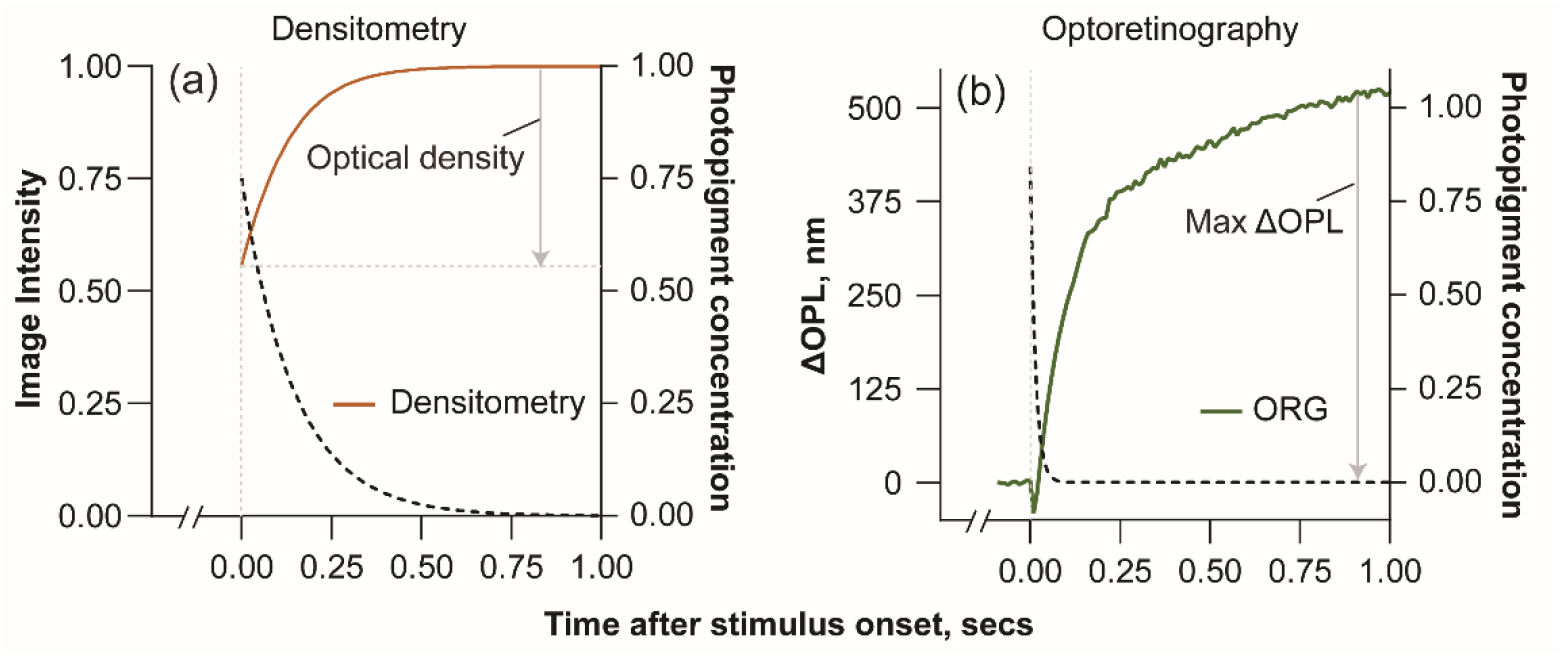
The dynamic range of ORG is ∼20x improved over densitometry. Densitometry (Fig. 1a) measures the time varying change in the backscattered image intensity (brown curve, left y-axis, Fig. 1a) from the retina as the photopigment concentration (black dashed curve, right y-axis, Fig. 1a) is reduced in the cone outer segment with a bleaching stimulus. ORG (Fig. 1b) measures the time varying change in ΔOPL in the cone outer segment (green curve, left y-axis, Fig. 1b) as the photopigment concentration (black dashed curve, right y-axis, Fig. 1b) is reduced with a bleaching stimulus. The dynamic range is the maximum obtainable change in the photopigment dependent signal attributed to a bleach (represented on the left y-axis for both), equal to ∼2 – 2.6x for densitometry and up to at least 50x for ORG.

In contrast, the stimulus evoked ΔOPL shows a significantly greater pigment-dependent change, scaling in magnitude from 10 – 500nm with increasing bleach from 0.3 - 50% [14]. Fig 1b shows ΔOPL vs. time for a 50% bleach using a 520 nm stimulus. Considering 10nm as the noise floor of the measurement, the ratio of post to pre bleach ΔOPL can be as high as 50x, indicating the dynamic range available to probe LMS-cone photopigments in ORG. The mechanism of action of ORG is currently unknown, though its bleach dependence suggests an involvement of the phototransduction amplification cascade[14, 19] that ultimately benefits its dynamic range.

In this article, we sought to investigate the correspondence between densitometry and ORG for cone classification in the same subjects and compare their fidelity for the task. First, we establish our ORG protocol for this task and ascertain its test-retest reliability by measuring the same subjects and eccentricities across different days. Variations in reflectance of cones caused due to interference, disc shedding and renewal, and shadows of blood vessels negatively affect both paradigms for classification [20-22]. Next, we ascertain whether cones with abnormal reflectivity exhibit normal function in ORG sufficient to identify their type. Finally, we compare the spectral assignments between the two paradigms with respect to the specific cone types and their spatial arrangement.

## 2. Methods

### 2.1 Reflective mirror based line scan AO-OCT

A line scan spectral domain OCT system equipped with AO, described previously [17], was used for ORG with cellular resolution. Briefly, a wavelength band at 820 ± 40 nm was selected from a supercontinuum light source (SuperK EXR-9OCT, NKT Photonics, A/S, Birkerød, Denmark) for imaging the retina. A 980±10 nm superluminescent diode (SLD) (IPSDD0906, Inphenix, USA) was used for wavefront sensing and AO correction. A 660±5 nm light emitting diode (LED) was used as the bleaching stimulus to differentiate the cone types because L-cones have 13 times higher sensitivity than M-cones, and S-cones have negligible sensitivity at this wavelength[23]. The wavefront sensing and the imaging beams were combined with a short pass filter and these two beams were combined with the bleaching stimulus using a long pass filter. Light backscattered from the retina was split into two beams using the same short pass filter in the detection arm. The 820 nm light scattered from retina was captured either with a line scan camera to optimize photoreceptor image focus in an *en face* image stream, or with a custom-built spectrometer for OCT. The spectrometer consisted of a diffraction grating (WP-600/840-35×45, Wasatch Photonics, USA) and a fast 2D camera (FASTCAM NOVA S16, Photron, USA). The fast 2D camera was used to capture the spatial and spectral dimensions along its 2 axes simultaneously to yield a cross-sectional OCT B-scan in a single frame. An anamorphic configuration was used to optimize the spatial and spectral dimension independently[16].

### 2.2 Subjects

Five color normal subjects were recruited for the study after an informed consent explaining the nature and possible consequences of the study. The research was approved by the University of Washington institutional review board and experiments were performed in accordance with the tenets of the Declaration of Helsinki. Of the 5 subjects, 1 was Caucasian male, two were female Caucasians and two were Asian males. Two of the color normal retinae were cone-typed earlier using densitometry, published in Sabesan et al. [11]. Subjects S3 and S4 in Sabesan et. al. are S4 and S5 respectively in this paper. In this study, cone classification for S4 and S5 with ORG, at 1.5 and 2 deg. eccentricity respectively was compared against data from densitometry in Sabesan et al., separated in time by 6-7 years. In all other subjects, cone classification was performed at 1.5 deg. temporal eccentricity. For comparison of cone classification across two days, subjects S2 & S4 were imaged at 4 deg. and 1.5 deg. eccentricity, respectively.

### 2.3 Imaging protocol

Subjects were dilated and cyclopleged using 1% tropicamide and/or 2% phenylephrine. They were placed in a dental impression to stabilize head movement. Their pupil was aligned to the optical axis of the instrument using a three axis translation stage assisted with a pupil camera. The aberrations were measured and corrected in real time using the wavefront sensor and deformable mirror. The retinal image captured in the line scan camera was optimized to focus the cone photoreceptors. The system was then switched to line scan OCT mode for further experiments. For ORGs, AO-OCT volumes were recorded after subjects were dark-adapted for 1-3 minutes to allow for sufficient regeneration of cone photopigments. The 660 nm, 20ms LED flash was delivered after the 10^th^ volume. The B-scan rate was 12000 scans/sec. Each volume contained 600 B-scans, and each video contained 50 volumes. Typically, unless otherwise stated, ten recordings were taken for averaging and improving signal-to-noise ratio. The schematic of imaging and processing is shown in Fig. 2 & 3.

**Figure 2:**
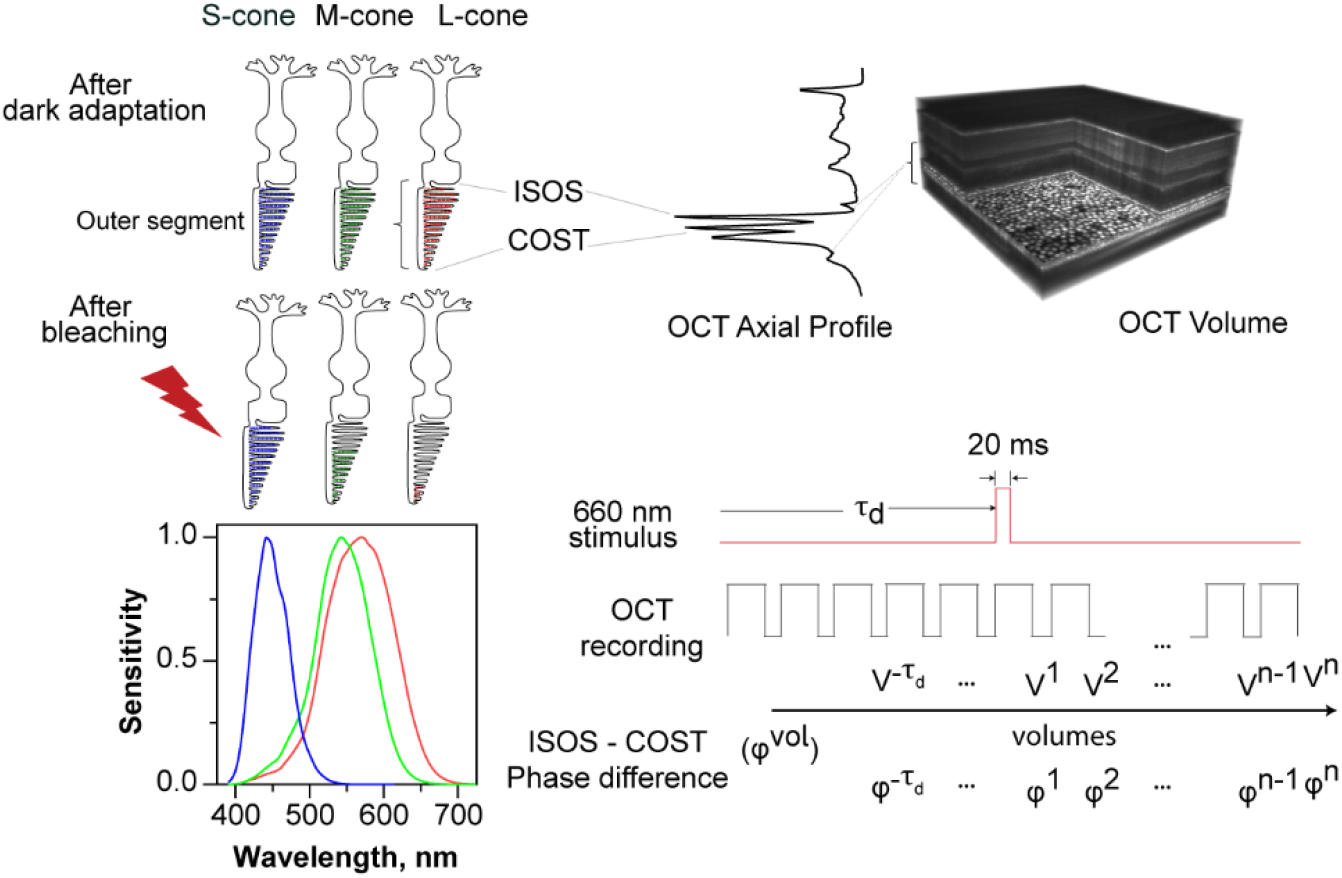
Schematic for cone classification using ORG. The registered AO line-scan OCT volume was segmented at the cone inner-outer segment junction (ISOS) and the cone outer segment tips (COST) to access the outer segment. After dark adaptation, photopigments regenerate in all cones. After a 660nm stimulus, L-cones have highest bleach & activity, followed by M-cones and then S-cones, as dictated by their decreasing spectral sensitivity to 660nm wavelength[23]. The onset of the 20 ms stimulus flash occurs at τ_d_ = 0.5 sec or after the 10^th^ volume. OCT volumes (n= 50) were recorded, reconstructed and registered. The phase difference between the ISOS and COST in each volume was calculated and treated to the steps in Section 2.4 to obtain a measure of OPL change in each cone, evoked by the stimulus.

**Figure 3:**
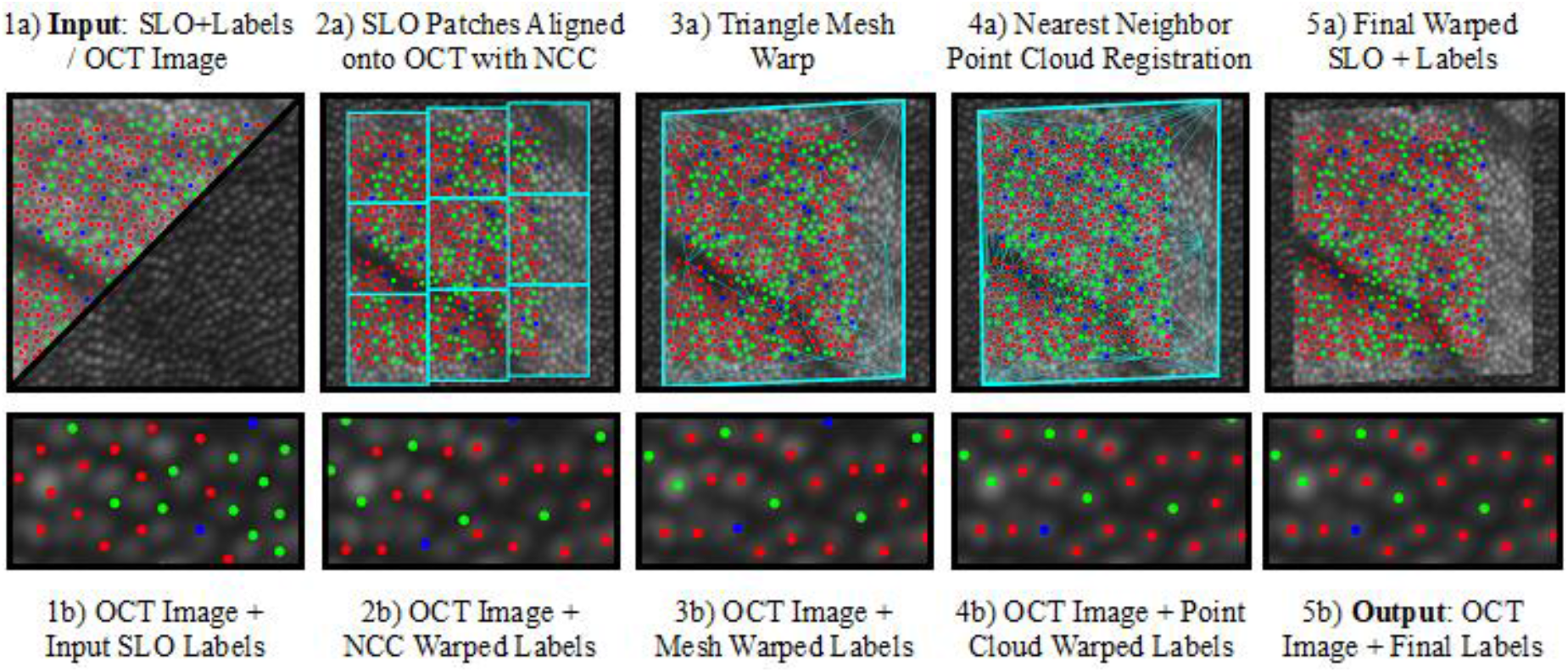
Transferring cone labels from an SLO image to an OCT image is a sequence of simple steps with the Retina Map Alignment Tool. These steps are shown left-to-right going from inputs to outputs. The LMS cone label types are shown as false-color RGB dots, and the SLO image is shown brighter than the OCT image for clarity. The bottom row shows a zoom-in of the SLO labels as they are gradually warped to align with the cones visible in the OCT image. From left to right: 1a) The inputs are the cone-labeled SLO image and the destination OCT image. 1b) Naively copying the SLO label positions directly onto the OCT image fails to pair the labels with the visible cones. 2a) We first rigidly match labeled patches of the SLO image to get a rough alignment with the OCT image. 2b) Notice how many SLO labels fall in the gaps between the cones, making pairings unclear. 3a) To improve alignment further, the patches are converted into a flexible triangle mesh, which the user adjusts in real-time (See Section 2.8 text). 3b) The labels are now unambiguously paired with the visible cones, but their positions are each offset slightly. 4a) To fix this, we find nearest neighbors between the two cone mosaics and produce new correspondences that snap each SLO label to the nearest OCT cone. 4b) Now every label is perfectly centered with each cone. 5a+b) The tool outputs the final SLO labels aligned to the OCT image, along with the warped SLO image. The user can now visually verify the transfer by checking cone-alignment between the warped SLO image and the destination OCT image.

### 2.4 OCT image processing

Typical steps were followed for OCT image processing. The details of the processing and ORG extraction are detailed in reference [16]. Each recorded 2D spectrum (*I*(*x, y, λ*)) was treated to background subtraction, k-space resampling (*I*(*x, y, k*)), and a Fourier transform *I*(*x, y, z*) to yield an OCT volume. All volumes were registered using segmentation-based 3D registration[24]. Once the volumes were registered, each OCT volume was referenced to the mean of all the volumes that were recorded before the start of the stimulus to cancel the arbitrary phase at each pixel. Then, the mean of 3 axial pixel complex values, centered at the boundaries of the ISOS, *I*_*ISOS*_ (*x, y*) and the cone outer segment tips (COST), *I*_*COST*_ (*x, y*) was calculated. The phase difference *I*_*ISOS*/*COST*_ (*x, y*) between these two layers was obtained by multiplying the complex conjugate of COST with the ISOS layer. The complex numbers *I*_*ISOS*/*COST*_ (*x, y*) were averaged over the collection aperture of a cone photoreceptor to yield one complex number per cone in a volume. The same was repeated for the 50 volumes in the 20 Hz time series to give *I*_*ISOS*/*COST*_ (*t*).

### 2.5 Complex average vs Angle average

We evaluated two approaches for averaging across repeat measurements to improve signal-to-noise. The first approach involved taking the *complex average* of individual ΔOPL vs. time series as below:

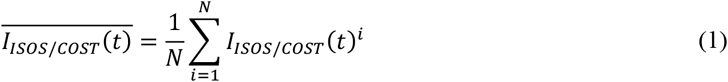

where N is the number of recordings, and *I*_*ISOS*/*COST*_ (*t*)^*i*^ is a complex valued time series for one measurement. The phase 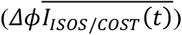 was computed by calculating the argument of the averaged complex time series. For phase responses that exceeded ±*π* radians, phase was unwrapped along the time dimension. The mean change in OPL 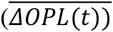 was calculated as

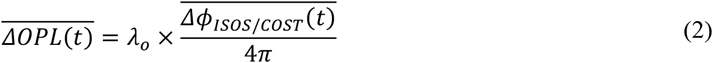

where *λ*_*0*_is the central wavelength equal to 820nm.

In the second approach, the argument of each individual complex valued time series was obtained to yield the phase, unwrapped if needed and converted to ΔOPL using equation 2. This approach represents the *angle average*. The mean 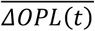 of all recordings was calculated as.

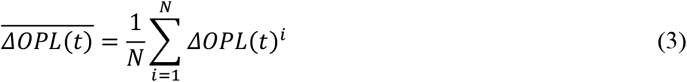

where N is the number of recordings *ΔOPL*(*t*)^*i*^ is the time series of ΔOPL for one measurement. An alternate strategy to the angle and complex average is performing the circular mean, a strategy used in cases where the data are inherently circular and/or periodic [25].

### 2.6 Clustering analysis

For segregating L, M and S cones, the ΔOPL averaged over the time after stimulus onset between 0.7 – 1 sec was calculated for every cone. A histogram of the time averaged ΔOPL (maximum ΔOPL elsewhere in this article) was fit with a sum of three 1-dimensional Gaussians. The clustering method followed here is analogous to a 3-component, 1-dimensional Gaussian mixture model[26]. The intersection of the component Gaussians separated the clusters into the three cone types. Each classified cone within a cluster is associated with an error percent or uncertainty, and a probability or likelihood of its assignment. Error is a group measure for S vs. M, and M vs. L-cones, defined as the area of overlap between their respective component Gaussians. Probability is a measure specific to each cone within a cluster, defined as the ratio of its component Gaussian value to the sum of all Gaussians.

### 2.7 Comparison of cone spectral classification across different days to assess test-retest reliability

To assess test-retest reliability, cone classification was conducted on two different days for two subjects, at one eccentricity each. Subjects S2 & S4 were classified at 1.5 deg. and 4 deg. eccentricity respectively. The recorded OCT volumes on both days were registered together to select the same cones for classification. Cones so classified were compared for cone-type specific mismatches between days. Bland-Altman analysis[27] was performed to compare the maximum ΔOPL across both days to evaluate their repeatability.

### 2.8 Cone-by-cone alignment for comparing densitometry & ORG

To compare outcomes of the two classification methods, we were first required to identify matching cones between the datasets obtained in AOSLO densitometry and AO-OCT based ORG. However, the two datasets obtained from these different instruments feature significantly different scales, orientations, cone appearances and motion-induced distortions. To address this problem, we used a specially designed Retina Map Alignment Tool. This tool aligns two cone-labeled retina images (‘retina maps’) via two-stages: automatic coarse image registration followed by human-assisted refinement and verification. This tool allows us to match hundreds to thousands of cones per subject with confidence in a fraction of the time it would take manually. See Figure 3 for a visualization of the pipeline.

The automatic alignment finds a piecewise linear warping between the images, which the operator can refine and verify through an interactive graphical user interface (GUI). The warping is parameterized by a sparse set of 2D pixel correspondence points between the SLO image and the OCT *en face* image at the COST. To obtain a continuous warping, a triangle mesh is constructed from the correspondence points. The tool then computes per-triangle transforms and applies texture mapping techniques to warp the SLO image onto the OCT image[28]. The initial set of correspondence points is found by first finding a rough translation, rotation, and scale between the images[29]. Then, the SLO image is split into 0.1 deg. wide patches and separately matched against the OCT image via normalized cross-correlation[30]. Finally, RANSAC is used to filter outlier matches[31]. The user verifies alignment by flipping between the two registered images and checking that each cone visually aligns with its match. If further adjustment is needed, the tool’s GUI allows the user to add, remove, or move these correspondence points and see the updated warp in real time. Once the user verifies the retina map registration, cone matches are identified by a nearest-neighbor point cloud registration. Individual matches can also be manually adjusted. Finally, the tool outputs a file with the cone coordinates and identities relative to the new image.

## 3. Results

### 3.1 Complex average vs Angle average

In this section, we compare the two methods of averaging – complex & angle average – across repeat measurements. Fig. 4a shows an example of a cone where the individual trial ORG traces have low inter-trial variability, low noise and no unwrapping artifacts. In this case, the angle average and the complex average traces align well with each other. Fig. 4b shows an example of a cone where repeat measurements are noisy and have unwrapping artifacts. These factors corrupt the angle average trace, and in this specific case, lead to a lower overall ΔOPL change compared to the complex average. This comparison was repeated for the same population of cones (n = 512) in Fig. 4c and 4d, where there is expected to be a mixture of cones with high and low inter-trial variability. The complex average of the constituent trials shows a clear separation between the three cone types in Fig. 4c. In comparison, the angle average contains more noise in the individual cone ΔOPL versus time traces leading to an ambiguity in the cone assignments. We follow the complex average for all data presented in this article. Phase variance is directly related to the inverse of signal-to-noise – higher the SNR, lower the phase noise and vice versa. The complex average effectively weights the phase information with the signal amplitude and is more robust to noise artifacts including phase wrapping.

**Figure 4:**
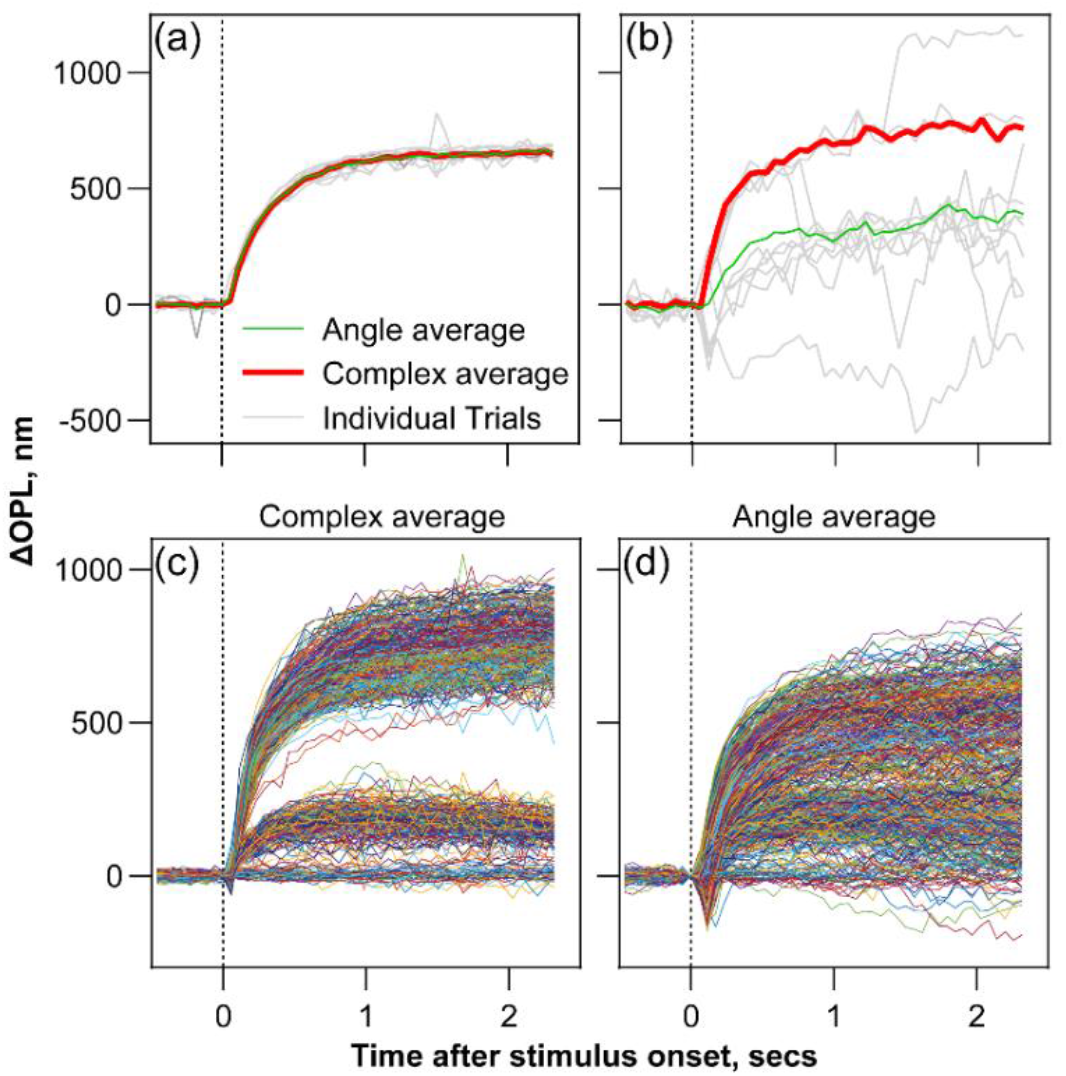
Comparison between complex and angle average for calculating the mean ΔOPL vs. time. Fig. 4a and 4b show examples for 10 repeat trials for 2 different cones in cases where the individual trials have low (a) and high (b) inter-trial variability respectively. When the two approaches are applied to a population of the same 512 cones, the complex average (Fig. 4c) readily segregates the cones into its respective spectral types. Angle average (Fig. 4d), on the other hand, is more susceptible to inter-trial variability and leads to ambiguity in the cone assignments.

### 3.2 Separating LMS cones

Figure 5 shows a representative example of the procedure for cone classification under a 660 nm stimulus bleach at 1.5 deg. temporal eccentricity in subject S4. The three cone types are well segregated owing to the variable bleach in each type. The absolute values of the registered AO-OCT complex volume segmented at the ISOS and COST (*I*_*ISOS*_ (*x, y*) and *I*_*COST*_ (*x, y*)) provides *en face* images revealing cone photoreceptor at the two layers (Fig. 5a and 5b). The mean ΔOPL between the ISOS and COST at time t = 0.5 sec after stimulus onset is shown in Fig. 5c for each cone (see Video 1 for time evolution of ΔOPL in individual cones). The magnitude of ΔOPL for each cone in the image is color coded according to the color bar, and shows the variability and distribution between cones. Three colors – red, yellow and blue on the color bar are dominant while the intervening colors are sparse, suggesting the correspondence to the three cone types. The time series of ΔOPL for individual cones is shown in Fig. 5d. The maximum ΔOPL for every cone is plotted as a histogram in Fig. 5e. The histogram is separated into the corresponding clusters of L, M and S cones by the intersection of the three component Gaussians. The individual cone ΔOPL traces are shaded red, green and blue in Fig. 5f according to their cluster assignment in Fig. 5e. The mean of the saturated ΔOPL for L, M, and S cones was 585±52 nm, 157±20, 9±18nm respectively. In each cluster, noisy ΔOPL traces and wrapping artifacts are filtered manually. Individual trials with excess eye movement or noise are discarded prior to the complex averaging. After these steps, a few cones exhibit ORG characteristics that are insufficient to unambiguously assign their type. These are excluded from the classification and labeled as ‘not classified (NC)’.

**Figure 5:**
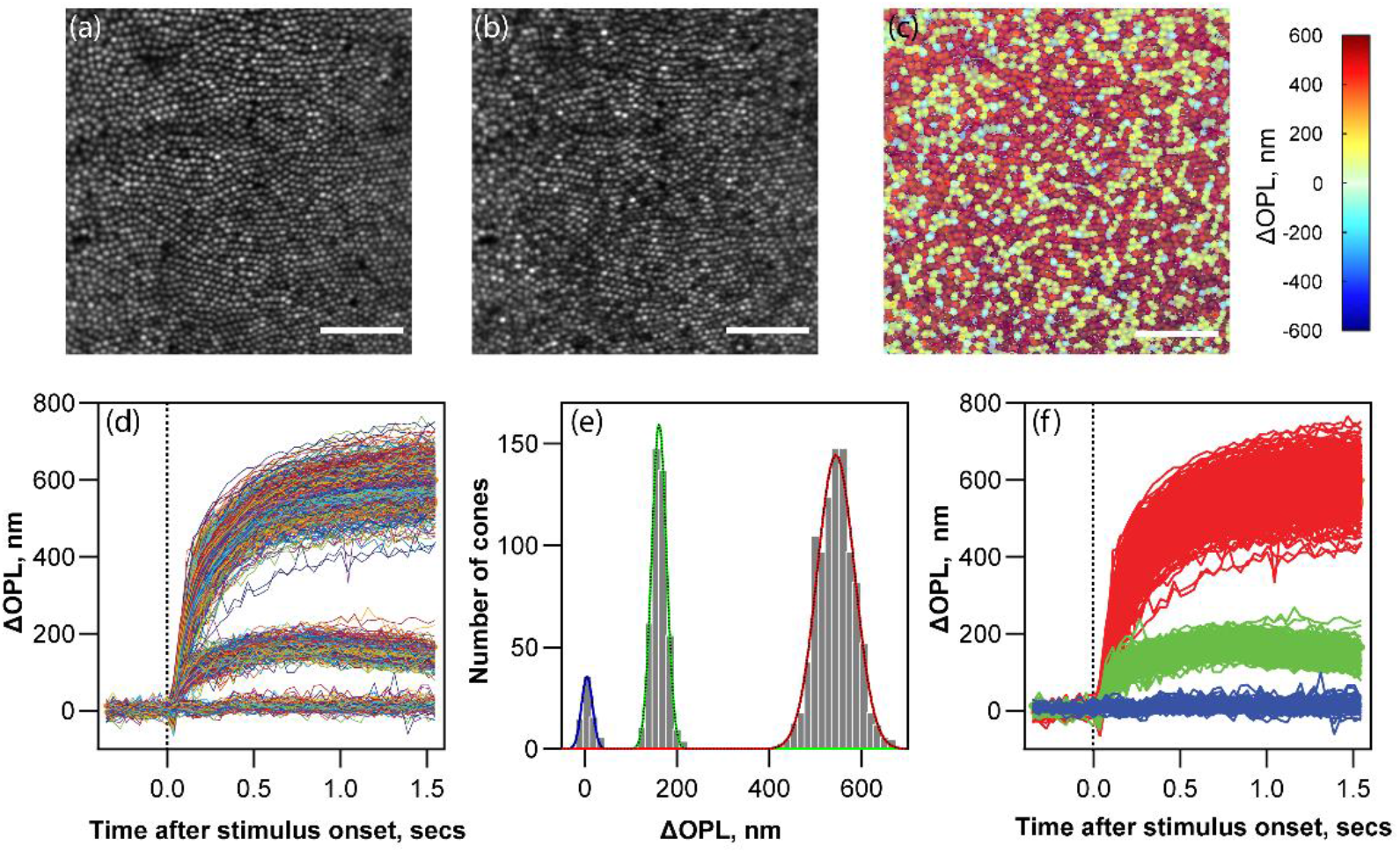
Representative example for cone classification for Subject S4. Registered AO-OCT volumes segmented at ISOS and COST yields the *en face* cone photoreceptor images at these layers in (a) & (b) respectively. The mean ΔOPL in cone outer segments at time t=0.5 is color coded according to the color bar in c) and overlaid on the cone mosaic in the COST image. Individual cone time series of ΔOPL is plotted in Fig. 5d. A histogram of the maximum ΔOPL is shown in Fig. 5e. Gaussian mixture model clustering analysis is used to segregate the cone types based on the maximum ΔOPL in the histogram. Fig. 5f shows the same traces from Fig. 5d, now color coded as red, green and blue to represent L, M and S-cones respectively. Scale bar: 10 arcmin

Figure 6 shows the cone classification for three subjects (S1, S2, & S3, from left to right columns) following the procedure in Fig. 5. Fig. 6 (a,b,c) shows the gray scale image and (d,e,f) shows the histogram of ΔOPL and cluster separations into the three cone types. Fig. 6 (g,h,i) shows the corresponding LMS-cone mosaics labeled in false-color according to their segregation obtained from Gaussian mixture model clustering analysis. Table 1 shows the total number of cones classified, percentage of S-cones, L:M ratio, the variation of saturated ΔOPL in each cone subtype, the probability and error of assignment for the 3 subjects. The average maximum ΔOPL between these 3 subjects for L, M, S-cones was 648±67, 170±28, 15±15nm respectively. The remaining two subjects appear below in section 3.5 In the 5 color normal subjects, the L:M cone ratio ranged between 1.5 - 2.4, and S-cone percent between 4.8 – 8.4 %, and both are comparable to the literature[9, 32].

**Figure 6:**
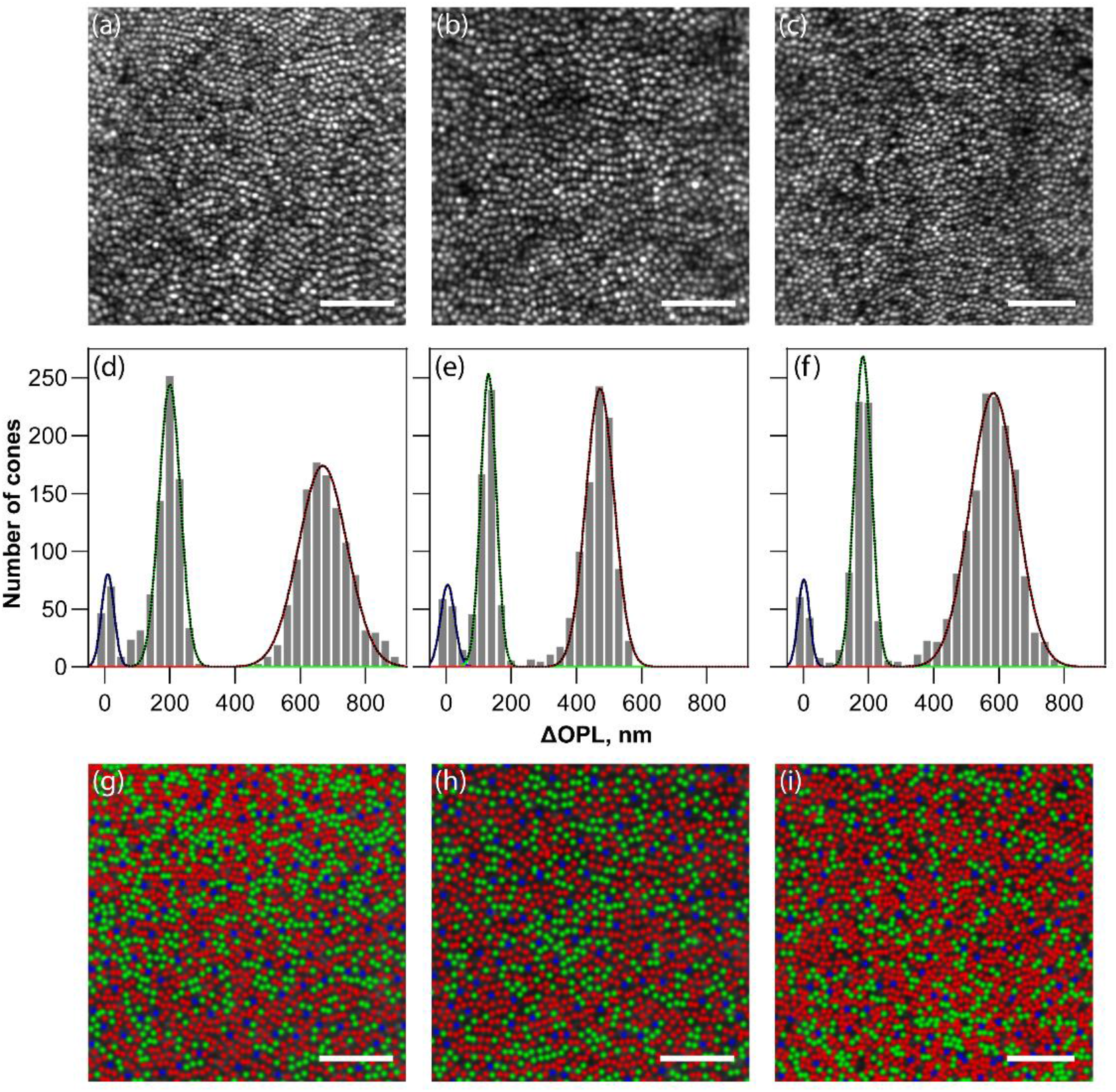
Cone spectral classification in three color normal subjects using ORG. Subjects S1, S2 & S3 are shown from left to right columns. (a-c) Gray scale *en face* images obtained by segmenting the registered and averaged AO-OCT volume at the COST (d-f) Histograms of maximum ΔOPL calculated in the time window 0.7 – 1 sec after stimulus onset overlaid with the component Gaussian fits obtained from Gaussian mixture model clustering analysis. (g-i) LMS-cone mosaics false-colored as red, green and blue to represent the three cone types. Scale bar: 10 arcmin

**Table 1.** Cone Classification Summary for 3 color normal subjects at 1.5 deg. eccentricity.

| Subject<br>/Age<br>/Ethnicity | # Cones | # Cones not<br>classified (NC) | L:M ratio | %S | Average Max $\Delta OPL$<br>(Mean ± Std) | | | Probability<br>(Mean ± Std) | %SM<br>error | %ML<br>error |
| --- | --- | --- | --- | --- | --- | --- | --- | --- | --- | --- |
|  |  |  |  |  | L-cones | M-cones | S-cones |  |  |  |
| S1/30/C | 1945 | 4 | 1.5 | 6.6% | $746 \pm 92$ | $187 \pm 41$ | $19 \pm 21$ | $0.99 \pm 0.04$ | 0.01 | $5e-4$ |
| S2/44/A | 1556 | 1 | 1.7 | 7.3% | $559 \pm 55$ | $130 \pm 28$ | $18 \pm 11$ | $0.99 \pm 0.02$ | 0.22 | $2e-5$ |
| S3/25/C | 2173 | 3 | 2.4 | 5.3% | $620 \pm 79$ | $177 \pm 30$ | $11 \pm 15$ | $0.99 \pm 0.03$ | $2e-3$ | $2e-3$ |

### 3.3 Dysflective cones

Cone photoreceptor reflections are variable, potentially due to a combination of interference of multiple reflections from the outer segment, disc shedding and renewal, abnormal waveguiding and perhaps other unknown factors[7, 8, 20, 21, 33, 34]. In addition, it is also known that a visible stimulus causes reflectivity variations that can be analyzed to infer their function, also termed intensity ORG, or iORG[35-37]. Support that these variations are caused due to interference from multiple reflections from the outer segment comes from Cooper et al.2020 [36]. They used a long coherence light source greater than the length of the outer segment that reduced the variability in cone reflectivity considerably. In the extreme case, some cones have abnormally low reflectivity, dubbed ‘dysflective’[38, 39]. It has been noted that normal eyes with no known disease have patches of dysflective cones that sometimes regain their reflectivity over time[40]. Also, using retinal tracking and targeted stimulus delivery, it has been shown cones with low reflectivity exhibit normal perceptual sensitivity[41]. In this section, we sought to investigate whether any potential functional deficit in ORG can be observed in dysflective cones.

Image formation in OCT relies on optical interference, and it may be plausible that reflections from some cones undergo destructive interference to exhibit abnormally low reflections. In the AOSLO, cones lying under the shadows of blood vessels have lower apparent reflectivity, especially when imaged with a visible wavelength as in retinal densitometry. Examples of dysflective cones are shown in Fig. 7 in subject S4 (a) and S5 (b) in their respective AO-OCT *en face* images of the COST. With AO-OCT alone, it was not possible to ascertain whether cones were indeed present in these dark areas indicated by the yellow circles. However, with the ‘Retina Map Alignment Tool’ described in 152 cl:801 section 2.8, we observed cone reflections present in the corresponding AOSLO image from the same subjects at these locations (S4: n=2 cones, S5: n=3). Fig. 7c and 7d show the ΔOPL versus time for the dysflective cones indicated in Fig. 7a and 7b after a 660nm stimulus bleach. For these dysflective cones, the phase difference was calculated at the same axial locations as where ISOS or COST would have been had there been a reflection from them. This axial location was determined based on the neighboring cones’ reflections. This approach assumes that the axial reflection locations don’t change substantially in a local area. These traces indicate normal ORG characteristics consistent with their classification into either L-cones and M-cones, based on their high and low maximum ΔOPL respectively. This indicates, that in addition to having normal perceptual sensitivity, dysflective cones exhibit normal functional responses in the ORG and transduce light with similar efficacy as cones with normal reflectivity.

**Figure 7:**
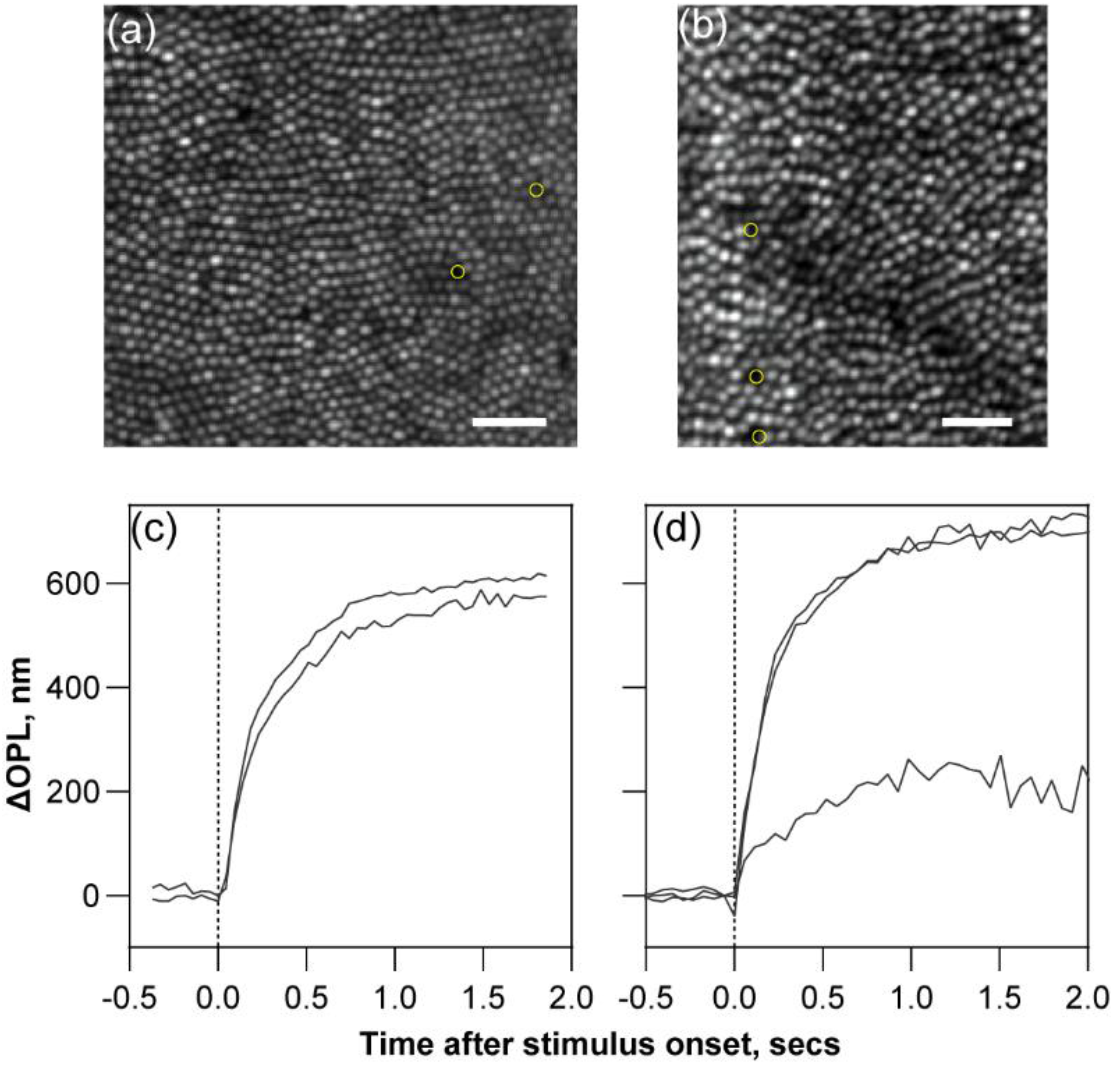
Optoretinography in dysflective cones. Fig. 7a & b show AO-OCT grayscale images for subject S4 (a) & S5 (b) at the COST layer where yellow circles indicate cones whose reflections were very low or absent. That cones were present in these ‘dark’ spaces was confirmed by finding their locations in the corresponding AOSLO image in the same subjects using the ‘Retinal Map Alignment Tool’ (see section 2.8). Fig. 7c & d show the corresponding ΔOPL vs. time traces showing normal characteristics consistent with their classification into L-cones (Fig. 7c) and L & M-cones (Fig. 7d). Scale bar: 5 arc-min.

### 3.4 Test-retest reliability in ORG cone classification

Figure 8 evaluates the test-retest reliability of classification across two different days. Fig. 8a) & c) show the maximum ΔOPL across the two days, labeled ‘day 1’ & ‘day 2’, at 1.5 deg. and 4 deg. eccentricity in Subjects S4 & S2 respectively. The maximum ΔOPLs obtained in each cone for the two days are plotted against one another; each data point represents an individual cone. Three distinct clusters with increasing maximum ΔOPL are observable corresponding to the L, M and S-cones. Cones were classified with high fidelity using Gaussian mixture model analysis based on each day’s measurements alone, with errors less than 0.02% and mean probability of 0.99. The distribution of data points in both x and y dimensions represents the variability in ΔOPL for each day, increasing in magnitude with the maximum ΔOPL. A few data points in each case lie outside the clusters and beyond the equality line. These correspond to the cones that did not match across days in their magnitude of maximum ΔOPL. Phase wrapping errors that lead to 2π phase ambiguities can partly explain these mismatches where a few cones exhibit high maximum ΔOPL on one day and low on the other. The mean Euclidean distance from the origin for the mismatched cones was 340 nm; in comparison, for the OCT center wavelength of 820 nm, a 2π ambiguity would result in a 410 nm error in ΔOPL.

**Figure 8:**
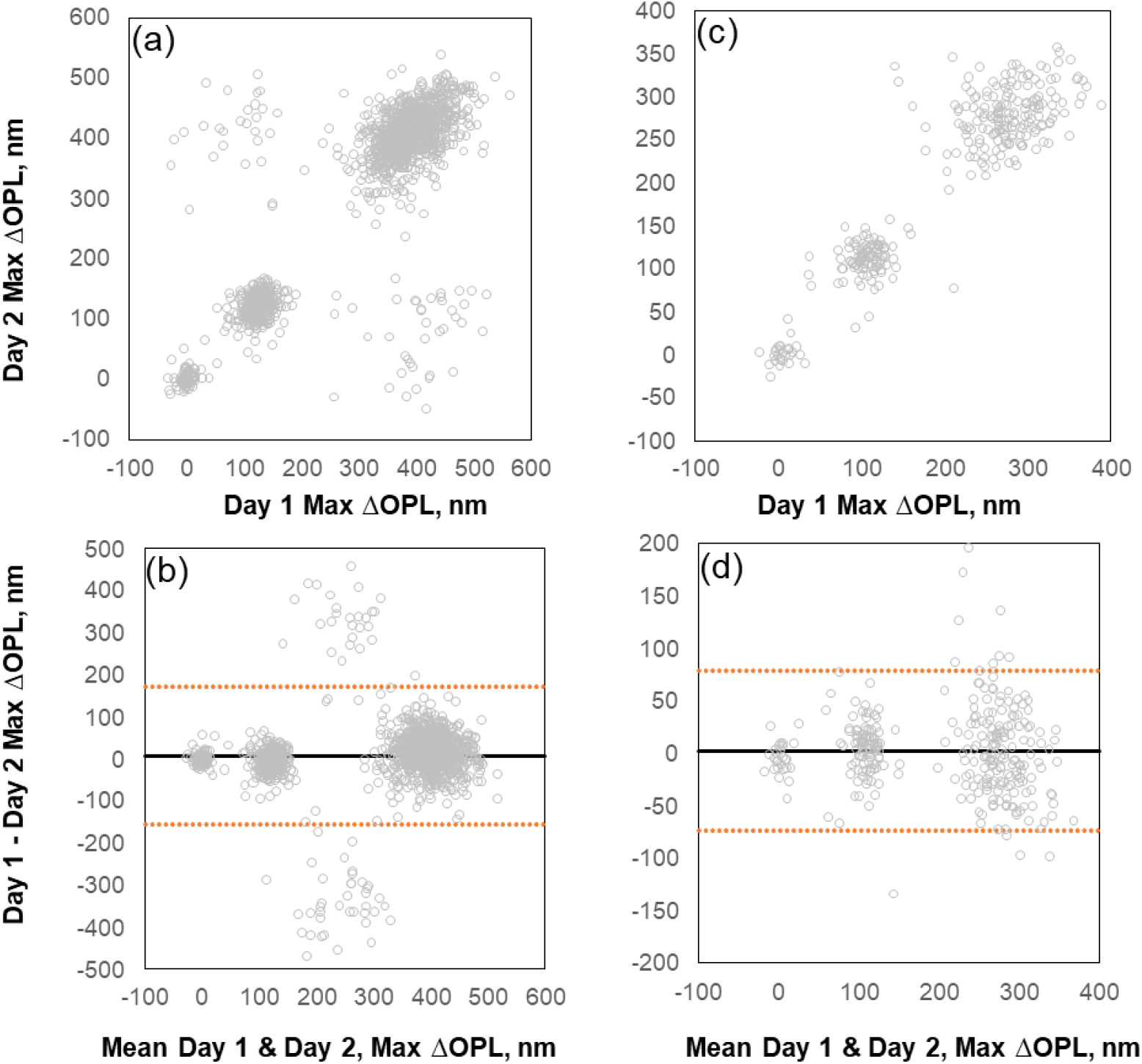
Comparison of maximum ΔOPL across two days, labeled day 1 & day 2, in the same cones. Fig. 8ab) & cd) show data at 1.5 and 4 deg. eccentricity for S4 & S2 respectively. Note the different y-axis scale at the two eccentricities. In a) & c), the magnitude of maximum ΔOPL for days 1 and 2 is plotted against each other, b) & d) show Bland-Altman plots for difference vs. mean of the maximum ΔOPL across the days. Horizontal orange line indicates ±95% confidence intervals in the difference between days, and the horizontal black line indicates the mean difference or bias.

To analyze the agreement in maximum ΔOPL as a whole between days, Fig. 8b) & d) show a Bland-Altman plot for 1.5 deg. and 4 deg. eccentricity respectively. Each data point is the measurement from the same cone across days. The three clusters corresponding to the cone types & the greater magnitudes of variability with increasing maximum ΔOPL are also observed in these plots (also see the standard deviation for S, M and L-cones in Tables 1 and 2). Typically, Bland-Altman plots represent the relationship between the difference (y-axis) versus the mean (x-axis) of two measurements to gauge their agreement. At 1.5 and 4 deg. eccentricity, 65 out of a total of 1397 cones and 13 out of a total of 305 cones lie beyond the ±95% confidence interval (orange line) respectively. The bias (black line), equal to the mean difference across all cones for both days, is meant to indicate any potential systematic difference between the days. The values of bias were 8.06 & 2.45 nm for 1.5 and 4 deg. respectively, and is close to the noise floor of the measurement for the ORG paradigm in the line-scan OCT [16]. At 1.5 deg. eccentricity, of the 1397 cones classified across both days based on Gaussian mixture model analysis on maximum ΔOPL, 4.2 % (59 cones – 48 L/M, 9 L/S, 2 M/S) were inconsistent in their assignment between the days. At 4 deg. eccentricity, 305 cones were classified across both days, of which 2.6% (8 cones – 3 L/M, 0 L/S, 5 M/S) were inconsistent in their assignment between days. Overall, a high agreement was noted between assignments across different days. The slightly larger inconsistency at 1.5 deg. could be related to phase wrapping errors. It is likely these were not sufficiently eliminated due to inadequate repeat measurements - 6 repeats on day 1 & 5 repeats on day 2 at 1.5 deg. - compared to the 10 repeats on each day at 4 deg. Another potential reason could be the higher cone density and smaller cone sizes at the closer eccentricity that may pose greater noise in the measurements across days.

**Table 2:**
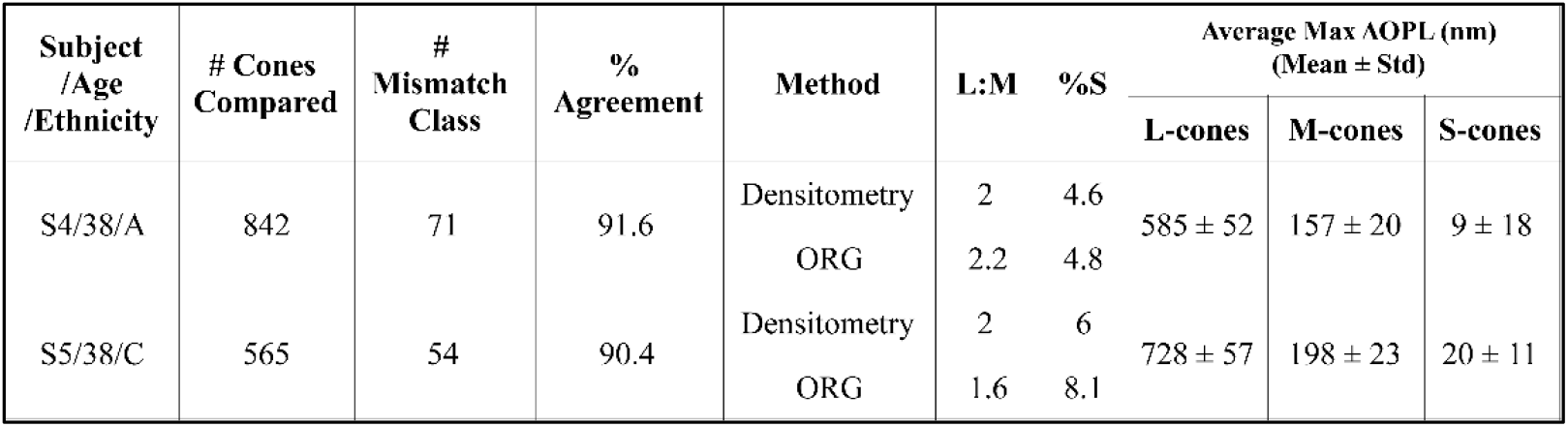
Summary of comparison between densitometry and ORG for cone classification.

### 3.5 Densitometry vs ORG classification comparison

We observed high classification agreement using densitometry and ORG ∼6-7 years apart in Subjects S4 & S5 in Fig. 9 and 10 (S4: 771/842 cones, 91.6%; S5: 511/565 cones, 90.4%). Fig. 9 shows the false-color LMS-cone mosaics from S4 and S5, classified at the same retinal eccentricity in densitometry and ORG. After cone matching and visual inspection by 2 separate graders, a few cones (S4: 12; S5: 4) present in Figure 9 were misaligned to a degree not amenable for comparison, and these were removed from further comparative analysis. Regions classified with ORG fully encapsulated regions classified with densitometry, and all cones classified with densitometry (S4: n=827, S5: n=559) were classified with ORG. If the comparison were restricted only to cones that were successfully classified in densitometry, the proportion agreement in S4 and S5 were 93.2% (771/827 cones) & 91.4 % (511/559).

**Figure 9:**
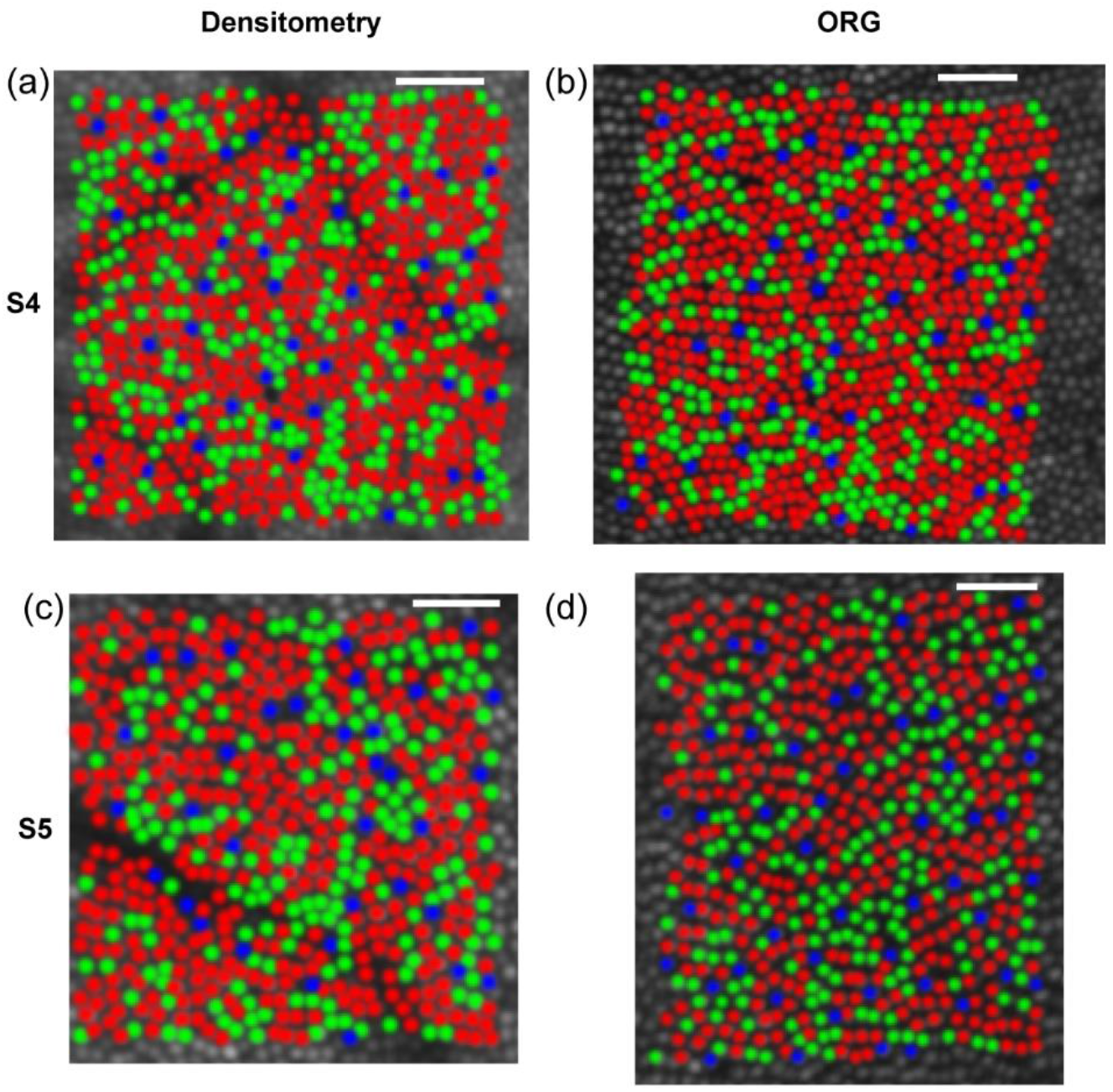
Cone spectral classification with densitometry and ORG. Fig. 9a & b show the cone spectral assignments in false color in S4 for densitometry and ORG, respectively overlaid on a grayscale image. Fig. 9c & d show the same for S5. Scale bar: 5 arc-min.

**Figure 10:**
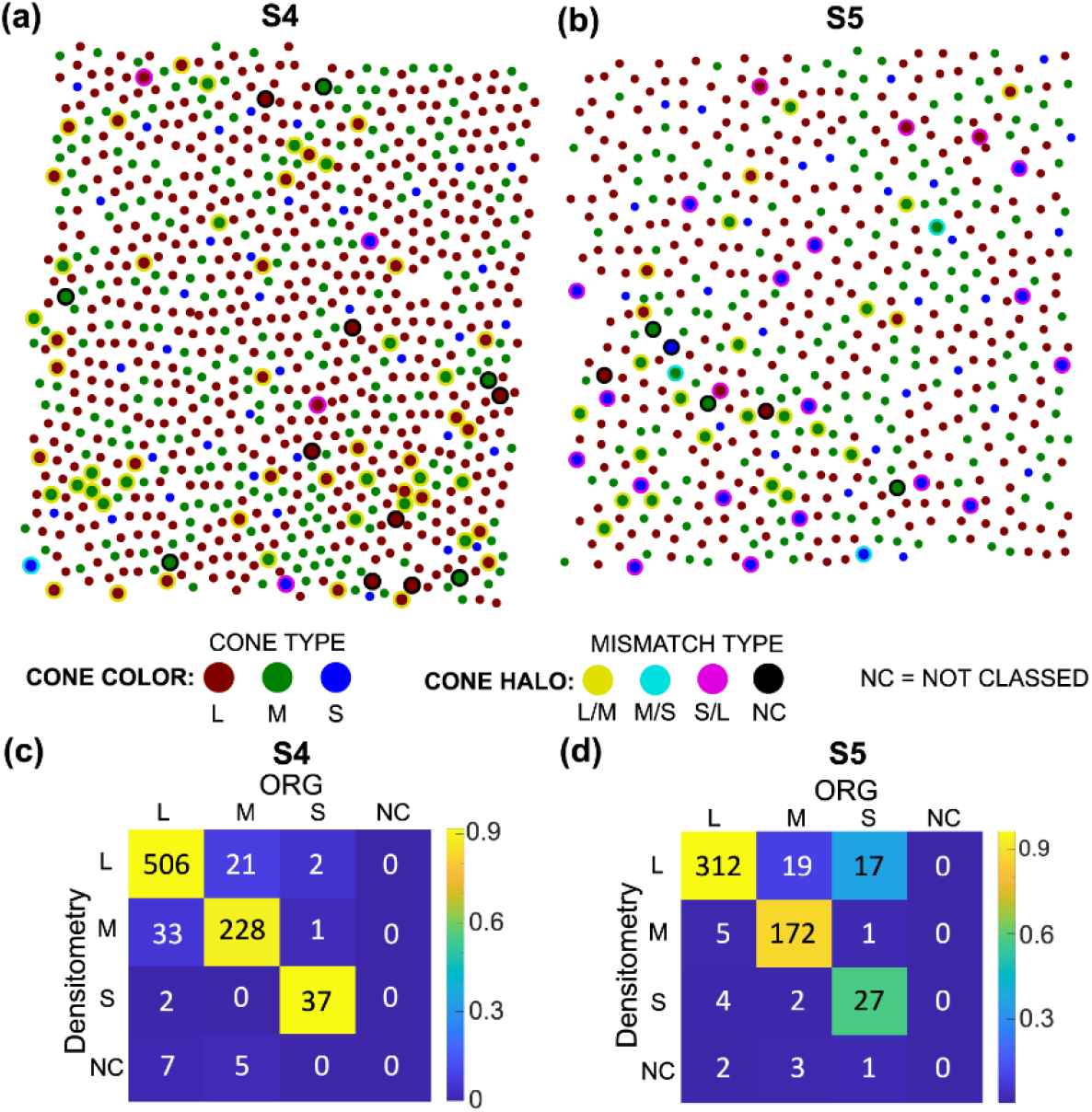
Classification mismatch between densitometry and ORG. Fig. 10a) & b) show ORG-classified mosaics highlighting densitometry-mismatches. Cone color indicates ORG LMS-classification. Cones with mismatched classification are plotted larger with a halo that indicates their densitometry classification. Fig. 10c) & d) show classification confusion matrices which quantify the number of cone-type specific matches in ORG and densitometry. Colors of each square and the corresponding color bars indicate the proportion matched between both modalities. Fig 10ac) are S4 classification mismatch and confusion matrix. Fig 10bd) are S5 classification mismatch and confusion matrix.

A small number of cones were not visible in the AO-OCT image (dubbed dysflective cones as section 3.3 above), but were nonetheless classified thanks to their transferred location from the densitometry image (see ‘Retina Map Alignment Tool’ in section 2.8 and Figure 7) (S4: n=2, S5: n=3). Finally, several cones which were not classified (NC) with densitometry owing to low reflections, in part caused by their lying under shadows of blood vessels, were classified with ORG (S4: n=15, S5: n=6).

A cone-by-cone summary of classification agreement across the two paradigms is summarized in Fig. 10 (a-d). Fig. 10a & b shows the distribution of cones across the regions of interest and how their cone assignment is consistent or different between the two modalities in S4 and S5. The background cone assignments are obtained from the ORG. Comparing the assignments as a whole in this manner helps appreciate potential factors related to the spatial arrangement or specific cone-types that may lead to inconsistencies. For example, it has been suggested that optical blur in densitometry might lead to an error in classification where cones of like-type are clumped together[4, 9]. Visual inspection shows few examples in both subjects where patches of 3-6 contiguous cones are mismatched. Two additional features are worth noting. Cones lying near the edge of the region-of-interest have a higher likelihood for mismatch than the center. One potential reason for this is eye motion that reduces the number of samples averaged at the borders compared to the center. Second, the bottom left quadrant of the region-of-interest in subject S5 has higher degree of mismatch, a location that has blood vessels overlying the cones casting a shadow, as shown in the gray scale image in Fig. 7b. This causes decreased back reflectivity of cones lying in the shadow, increased absorption by the blood vessels for the bleaching light and increased forward & backward scatter of the imaging light in both modalities.

Confusion matrices in Fig. 10c & 10d summarize the mismatch by cone type in both modalities. In both subjects, classification agreement (relative to ORG assignment) was highest for L-cones (S4: 92%, S5: 97%), decreasing for M-cones (S4: 90%, S5: 88%) and S-cones (S4: 93%, S5=59%). Table 2 summarizes the classification agreement, L:M ratio & % S-cones measured from each method. Mismatches had a slight effect on the L:M ratio measured for both subjects (S4: 2.0 vs 2.2, S5: 2.0 vs 1.6), negligible effect on the percent S-cones measured for Subject S4 (4.6 v 4.8%), and a more substantial effect on percent S-cones measured for Subject S5 (6.0 vs. 8.1%). For Subject S4, classification mismatches were dominated by L/M mismatches (of 71 mismatches, L/M = 76.1%, S/(L&M) = 7.0%, NC = 16.9%), whereas a higher proportion of mismatches for subject S5 involve S-cones (of 54 mismatches, L/M = 44.4%, S/(L&M) = 44.4%, NC = 11.1%). In particular, for Subject S5, a disproportionate number of cones assigned S in ORG were assigned L with densitometry (17/54 mismatches). Of these 17 L vs. S-cone mismatches, 5 occur at the borders of the region-of-interest, and 7 occur in the area overshadowed by the blood vessel (bottom left quadrant of Fig. 10b).

## 4. Discussion

This article presents the optimization and characterization of cone spectral classification using ORG and its comparison against densitometry. The cone mosaic of five color normal subjects were classified with ORG, showing high probability (∼0.99), low error (<0.22%), high test-retest reliability (∼97%) and short imaging durations (< 1 hour). The values for error are comparable to those reported previously for cone classification with point-scan AO-OCT[13]. The L:M cone ratio, and percent S-cones measured in this cohort are consistent with literature[9, 32]. In comparison, slightly lower probability (∼0.95) and higher error (∼ 5%) was observed in densitometry for classifying cones[11]. Further, it was suggested that the imaging sessions be distributed across a few days to account for reflectivity variations in cones, independent of photopigment absorption, leading generally to a greater time needed per subject[9]. Averaging is essential for both modalities given how ORG cone assignments can also be affected by phase wrapping errors when there are inadequate repeat measurements.

The overall higher fidelity of ORG is the result of greater than 20-fold higher dynamic range available in ORG compared to densitometry for capturing the perturbations caused by bleaching in the backscattered reflection from cones. Since the OCT layer reflections encasing the outer segment are used for computing the light-evoked ΔOPL, the optical signal is largely confined to the cone outer segment and not corrupted by reflections from other layers. Optical phase difference used to compute the ΔOPL has high sensitivity of about a few nanometers when phase is referenced between the ISOS and COST to remove common-mode noise. The bleach dependence of the ΔOPL is non-linear, scaling in magnitude with a gain greater than 1 suggesting a relationship to the amplification cascade of phototransduction[14]. The mechanism of the change in outer segment OPL has been attributed to the influx of water to maintain osmotic balance during phototransduction[19]. This is suggested to be in response to the disturbance of osmotic balance caused by the intermediary osmolytic products of the cascade that are amplified in their concentration compared to photopigments. While this multiplicative effect provides a gauge for photopigment concentration sufficient to classify cones with high fidelity in healthy eyes, the intermediary osmolytes are currently unknown. This unknown is critical to recognize when using ΔOPL as a biomarker for retinal diseases where mutations in specific proteins of phototransduction are implicated. In such cases, ΔOPL will not directly provide a gauge for photopigment concentration.

These aforementioned benefits of ORG do not apply to densitometry. Since absorption imaging is not depth-resolved to cone outer segments, it is confounded by the contribution of stray light due to scattering from other retinal layers, blood flow and anterior optics. With the confocality provided by AOSLO, the contribution of stray light can be reduced substantially. Still, maximum measurable change in absorption is limited to 0.41 log units (2.6-fold) at 550 nm[18]. Changes in optical intensity due to absorption cannot capitalize on the benefits of phototransduction amplification. The variations in cone reflectivity following a stimulus, as observable in iORG, have been used to separate S-cones [36] and can in principle be used to infer all three types.

Given that the mechanism of densitometry is well established and attributed directly to photopigment concentration, it was essential to compare the cone assignments obtained by the ORG method against it. A high agreement (mean: 91%) was observed between the two modalities in 2 subjects, with measurements conducted 6-7 years apart. Cones lying in the shadows of blood vessels in AOSLO were likely to be inconsistent in their assignments. Contiguous clusters of 3-6 mismatched cones were observed, attributable to optical blur & scatter. A greater degree of mismatch in S-cones was observed, likely in part due to the fact that S-cones are identified by a lack of their response to the bleaching stimulus in both modalities. Instead, a bleach that activates S-cones could be used in the future. Dysflective cones were found in both modalities. These cones were located in their counterpart image, and analyzed for their spectral type using ORG. Such cones with abnormal reflections have been observed in healthy and diseased eyes[39, 40, 42]. They have previously been shown to have normal function also[41, 42]. That they exhibit normal functional responses in the ORG has important clinical implications. A preponderance of ‘dark’ or dysflective cones in pathological eyes may not provide the complete picture of the dysfunctional state, and a functional measure of phototransduction such as ORG may be a more sensitive biomarker.

In summary, the protocol for cone classification was characterized using ORG and compared against densitometry. The benefits obtained in ORG from its higher sensitivity and dynamic range led to high efficiency and fidelity for cone spectral classification.

## Acknowledgement and Disclosures

Burroughs Wellcome Fund (Careers at the Scientific Interfaces Award); Research to Prevent Blindness (Career Development Award, Unrestriced grant to UW Ophthalmology); National Eye Institute R01EY029710, P30EY001730, U01EY032055); Weill NeuroHub Next Great Ideas Program; Air Force Office of Scientific Research (FA9550-20-1-0195 and FA9550-21-1-0230).

VPP and RS have a commercial interest in a US patent describing the technology for the line-scan OCT for optoretinography

